# Addressing Data Fragmentation in Biodiversity: A Workflow for integrated Species Distribution Models

**DOI:** 10.64898/2026.05.19.721053

**Authors:** Sam Wenaas Perrin, Kwaku Peprah Adjei, Philip Stanley Mostert, Ron Ronald Togunov, Ivar Herfindal, Joachim Paul Töpper, Jon Arvid Grytnes, Joseph Chipperfield, Robert Brian O’Hara, Anders Gravbrøt Finstad

## Abstract

**Aim:** A comprehensive understanding of the spatial distribution of biodiversity is hindered by fragmented datasets, sampling biases, and inconsistent observation protocols. Here, we present a workflow that integrates disparate datasets to produce large scale maps of biodiversity metrics as a basis for management-relevant information tools. We use integrated species distribution modeling (iSDM) to account for sampling biases and disparate data collection techniques, taking advantage of the vast numbers of open datasets available in data aggregators like GBIF.

**Location:** Norway (excluding Svalbard and Jan Mayen)

**Taxon:** Vascular plants

**Methods:** The workflow consists of four main steps: data acquisition, data integration, integrated species distribution modelling (iSDM), and the production of derived outputs. Input data include structured surveys, opportunistic observations, and environmental covariates. These are standardised and integrated into a point-processed based iSDM framework to produce species richness maps, associated uncertainties, and sampling effort maps. The outputs are further processed to identify biodiversity hotspots or to summarise species–environment relationships.

The workflow used vascular plant data from Norway, combining occurrence-only and presence-absence datasets with environmental covariates. Outputs were generated at a spatial resolution of 500 x 500 meters, balancing accuracy, computational feasibility and relevance for management decisions. High-performance computing resources were utilized for model fitting and predictions. A subset of available data was used to validate the species richness maps.

**Results:** We produced detailed maps of species richness, uncertainties and sampling intensity across Norway’s heterogeneous landscape, incorporating 1218 species in our final results. The species richness patterns highlight patterns consistent with previous mapping efforts. Validation showed an increase in model accuracy when compared to models which did not use an iSDM framework. The workflow highlights limitations in the infrastructure of the currently openly accessible data, particularly the need for more structured presence-absence datasets and standardized metadata.

**Main conclusions:** This study underscores the potential of workflows that integrate disparate datasets for biodiversity modeling. To maximize accuracy and utility, future efforts should focus on improving data standardization, the publication and collection of more structured data, and fostering data-sharing collaborations. Advances in the workflow itself, including optimising modelling covariates and integrating more comprehensive spatio-temporal aspects, will also increase the relevance of the outputs. These advances will increase our ability to estimate species richness with a precision and accuracy that can reliably inform conservation and management decisions.

## Introduction

A comprehensive understanding of the spatial distribution of biodiversity is essential to effectively address contemporary challenges in ecology and conservation biology (Meyer, 2015). Most countries worldwide have committed to protecting and restoring natural areas of high biodiversity significance through the Kunming-Montreal Global Biodiversity Framework (Targets 2 and 3, Convention on Biological Diversity, 2022). However, without comprehensive knowledge of where we find key areas of high biodiversity significance, we can not reliably identify where protective measures and restoration efforts should be focussed, or assess their impact. This will likely result in fragmented and inefficient biodiversity management (Paloniemi et al., 2012; United Nations Environment Programme, 2021).

There are several major obstacles to providing this requisite knowledge. Even with a massive scaling up of citizen science efforts and new technology, biodiversity can not be physically mapped everywhere. A viable method to fill the resulting gaps is to develop predictive tools that can use the available data from sampled areas to estimate distributions of species in unsampled areas. Species distribution models (SDMs) address this challenge, leveraging the relationship between known species occurrences, environmental variables and the spatial patterns formed by these relationships to predict where species are likely to occur (Elith & Leathwick, 2009). A wide variety of SDMs have been developed, but the trustworthiness of each model relies on the quality and quantity of input data.

The development of large public biological databases such as the Global Biodiversity Information Facility (GBIF) has made large amounts of species occurrence data now openly available. This is produced from a diversity of sources, among them citizen scientists, traditional surveys, eDNA surveys and camera traps. This diversity increases the amount of available data, but the lack of standardization across these sources leads to heterogeneity in spatial, temporal and taxa-level bias. Much of the recent growth in data availability stems from citizen science data, which is particularly skewed towards areas that are more easily accessible (Bowler et al. 2022), and towards charismatic, well-known and easy to observe species (Barbato et al. 2021, Stoudt et al. 2022). These biases can skew model outputs if uncorrected, estimating higher biodiversity in areas that are more accessible (Qiao et al., 2015).

As a consequence of the wide range of environments and organisms being studied with a plethora of observation methods, the resultant variety of datasets presents substantial challenges for research synthesis. Model-based data integration is a powerful method for combining datasets while preserving their unique strengths (Isaac et al., 2020). Integrated Species Distribution Models (iSDMs) separate biological processes from sampling processes, enabling the integration of diverse data types collected through varying sampling methods. iSDMs provide a rigorous and flexible mathematical basis for reconciling and synthesizing different biodiversity datasets.

Because iSDMs can make efficient use of cross-species and cross-dataset information, they should be ideal tools for large-scale biodiversity assessments. Most workflows currently focus on analyses of individual datasets (whether presence/absence or presence only). Practical applications will require that many datasets and species be processed efficiently at the same time.To ensure operational applicability, the workflow must be standardised, requiring both species and environmental data to be formatted in a common, interoperable structure that enables a high degree of automation.

Here we present an open workflow for large-scale biodiversity assessment applying the iSDM modelling framework to many species in one workflow. The workflow imports a range of different datasets, standardises them to a common format, fits multi-species integrated distribution models, and produces a variety of data products that can be used by scientists and managers to understand patterns of biodiversity.

## Workflow considerations

Designing a data integration workflow which is robust, scalable, and suitable for reliable biodiversity analyses requires careful consideration of several key requirements:

### 1. Transparency and reproducibility

Ensuring transparency is critical, both to uphold scientific rigour and to support reproducible and accountable decision-making. This requires that all procedures, code and source data be openly accessible, supported by comprehensive documentation and appropriate licensing to facilitate reuse. The FAIR principles (Findable, Accessible, Interoperable, Reusable) (Wilkinson et al., 2016) should guide every stage of the process. Data must be actively shared through appropriate repositories (Tedersoo et al., 2021), ensuring that data provenance can be tracked. Tracking provenance ensures the trustworthiness of outputs and enables efficient revisions and updates as source datasets evolve.

### 2. Addressing biases in observations

Sampling biases remain a significant challenge in most contemporary datasets, often resulting in skewed predictions when uncorrected (Qiao et al. 2015; Elith & Leathwick, 2009). Unstructured presence-only datasets (henceforth referred to as “occurrence-only” data) often provide greater spatial and temporal coverage compared to more systematically collected data; however, due to the absence of detailed information about the observational process, sampling bias usually can not be inferred from these data (Yackulic et al., 2013; Pacifici et al., 2017). “Structured” datasets - henceforth used to referred to data which provides enough information on the sampling process to infer sampling events and sampling effort - require systematic collection under controls for effort, methodology, and spatial coverage. Such data are often limited in both spatial and taxonomic coverage (Hortal & Lobo 2005, Marta et al. 2019).

Methods combining strengths and weaknesses of different data sources have recently been shown to outperform models relying on a single data source. In these cases, the use of these models improved the accuracy of results and estimates compared to those based solely on individual data sources (e.g. Simmonds et al., 2020; Isaac et al. 2020).

### 3. Source Data Requirements

In species distribution modeling, data can be broadly categorized into two types: environmental covariates, which determine where species can exist, and biodiversity data, which document species occurrences. Species occurrence data relevant to species distribution modeling (SDM) can be broadly categorized into two types - the aforementioned occurrence-only and structured data.

The vast majority of openly available biodiversity data is registered as occurrence-only data, only yielding information about the species being detected at a given place and time. Detailed information on presence-absence observations or metadata on observation effort (e.g., observation method, sampling effort) is often omitted during data transcription and lost. While some of this information can be reconstructed, much of it is irretrievable.

Structured data must provide opportunities to infer information about the sampling process, enabling correction for biases. To allow estimation of likelihood of occurrence, such data should include records of the presence or absence of organisms during a given sampling event. While occurrence-only data covers a vast temporal, taxonomic, and spatial range, it is necessary to complement these data with structured datasets that enable the inference and correction of sampling bias.

### 4. Spatial Resolution

The choice of spatial resolution is particularly crucial for applying species distribution models (SDMs) in management. Fine-grained models are often preferred by practitioners, facilitated by increasingly detailed environmental datasets such as high-resolution land-cover maps. However, choice of resolution must go beyond the desire for increased detail (Meynard et al., 2023). First, alignment between the spatial resolution of covariates and species occurrence data is essential. Mismatches can bias estimates of species-environment relationships and undermine model robustness (Chauvier et al., 2022; Gaget et al., 2025; Mourguiart et al., 2024). Second, ecological processes influencing species distributions operate at varying spatial scales. At broader scales, distributions tend to be relatively stable (Lomolino et al., 2017; Gaston, 2023), whereas fine-scale occurrences are dynamic and stochastic (Hanski, 1999). Accurate fine-scale predictions often require disproportionately dense occurrence data to account for these dynamics. Third, spatial scale alters environment-occurrence relationships. Aggregating occurrences to a lower level of resolution can change the mean likelihood of occurrence and shifts optimal environmental values, which may enhance predictive ability if the scale better matches the species actual relationship with its environment (Meynard et al., 2023).

For models on a large geographical scale, additional challenges arise from geographic variability in environmental drivers and species composition, influenced by gradients in elevation, temperature, precipitation, and historical biogeography (McVicar et al., 2013). Spatial variability in observation density and ecological processes further complicates predictions, particularly for wide-ranging species with diverse ecological responses (Mourguiart et al., 2024). Although finer resolutions are intuitively expected to improve predictions, empirical evidence shows inconsistent results, with outcomes influenced by the interaction between spatial resolution and ecological complexity (Franklin et al., 2013; Meynard et al., 2023).

### 5. Computational Efficiency

Efficient computational workflows are essential for integration of the vast and complex information needed. Workflows must scale to handle relevant spatial extents, fine-grained environmental covariates, and diverse biodiversity data. This necessitates the use of optimized algorithms, parallel processing, and high-performance computing resources. There is therefore a considerable challenge in upscaling research-driven algorithms and statistical methods that are primarily tested and demonstrated using selected datasets and limited spatial and taxonomic scales.

Additionally, computational efficiency is critical for iterative model development, allowing researchers to refine parameters, test hypotheses, and produce results in a timely manner. A well-designed workflow should ensure that the trade-offs between speed, scalability, and model complexity are carefully balanced, making the approach practical for real-world applications in biodiversity conservation and management.

## Methods

### Methods overview and workflow at a glance

The workflow is divided into the processes of data acquisition, data integration, integrated Species Distribution Modelling (iSDM), and creating predictions and other derived products (Figure 1). The starting point of the workflow is a series of datasets categorised into three main classes; Structured data (Figure 1a), opportunistically collected data (figure 1b) and data on environmental covariates (Figure 1c). Each dataset is examined using the available metadata and subsequently restructured in order to ensure that the results from different models are able to be entered into a common integrated species distribution modelling framework (Figure 1d, described in Mostert et al. 2024a). The iSDM framework gives three primary results; maps of individual species intensities (Figure 1e), maps of uncertainties in the individual species intensity estimations (Figure 1f), and maps of sampling intensity pooled for species groups (Figure 1g). These then may be processed further, e.g. maps of hotspots of biodiversity can be drawn (Figure 1h), or analyses of the effects of covariates across species groups can be extracted. The analysis is then validated using real observed data from the study area (Figure 1i).

**Figure 1:**
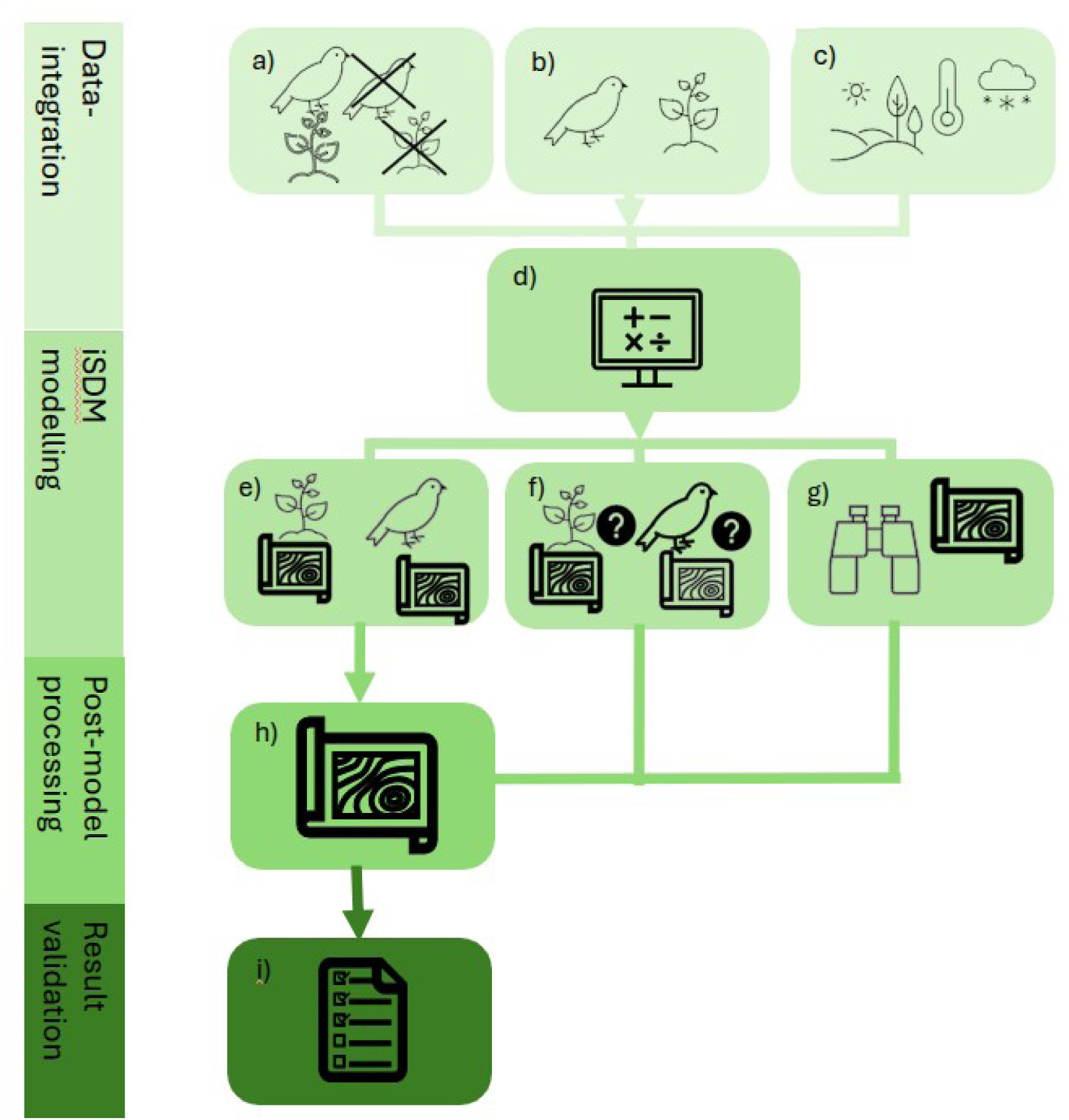
Visual summary of workflow which employs integrated Species Distribution models to produce species biodiversity metrics

Here we demonstrate the applicability of the workflow by producing maps for Norwegian vascular plants demonstrating species richness, uncertainty and sampling intensity. Vascular plants were chosen for demonstration purposes due to their fairly well known taxonomic status and their ample data availability from research projects, citizen science projects, natural history collections and structured monitoring.

All code used for the downloading, processing, modelling of data and subsequent production of species richness and sampling bias metrics is available at GitHub (Anonymous, 2026).

### Species occurrence data acquisition and processing

Species occurrence data can be obtained from a variety of sources. At a global scale GBIF (GBIF: The Global Biodiversity Information Facility 2024) provides FAIR access to an unparalleled source of biodiversity data, meeting criteria for openness and provenance tracking through a well developed citation system (GBIF.org 2024a). We downloaded species occurrence data using the GBIF API (GBIF.org 2024b) through the *rgbif* package (Chamberlain et al., 2023) in R (R Core Team 2023). We also filtered the occurrence records geographically by a multipolygon geometry depicting a simplified outline of mainland Norway, and also excluding any occurrence records having a coordinate uncertainty of more than 250 metres or a collection date before 2012 (GBIF.org 24 February 2026). We subsequently removed occurrences that were not identified down to species level, and all occurrences that were flagged by GBIF as having coordinates that were invalid or out of range.

Despite the success of standardization and semantic interoperability provided by Darwin Core (Wieczorek et al. 2012), GBIF only provides data as occurrence only. This means that after downloading the data directly from GBIF, post-processing and inference of content are needed in order to recreate and standardise information about the sampling process.

The downloaded datasets were examined and pre-classified into two groups based on their available metadata - occurrence-only and presence-absence (or, more precisely, observed/not observed). If the metadata provided insufficient information to decide, the listed metadata author was contacted and the necessary information was ascertained. If the necessary information was still not ascertained after this consultation, the dataset was left as an occurrence-only dataset. Datasets successfully classified as presence-absence were then downloaded and processed directly from IPT endpoints at which the unfiltered datasets are stored in DarwinCore Archive format in order to produce absences as required. Due to the labour intensive nature of this part of the workflow, we put a cap on the number of datasets included. After more than 98% of the records in the data download were classified, the remaining were treated as occurrence only.

At this stage, datasets were also assessed based on whether or not they were suitably area representative. Spatially representative presence-absence data covering the modelled area is required to ensure that the conclusions from our models are not significantly affected by sampling bias to certain areas (Boyd et. al. 2022). The GBIF download did not contain any representative data that could be classified as presence-absence. We therefore added open data from the Norwegian Environmental Agencies (“Area representative nature monitoring” - Arealrepresentativ Naturovervåkning - ANO) (Norwegian Environment Agency, 2025), which was reformatted and standardised to be interoperable with the GBIF data.

Lastly, any species which had less than 50 presence observations across all datasets were excluded from analysis.

### Environmental data acquisition

An initial list of environmental covariates was made in order to cover the majority of drivers of the distribution of species, while keeping the total number of covariates to a minimum in order to ensure comparability and reduce model processing time (Table 1). Interviews with two domain experts were carried out in order to identify covariates that would have an impact on the distribution of vascular plants. Scientists qualified as experts if they had experience working with vascular plants in Norway on large spatial scales. A full list of model covariates is available in supplementary table S1.1. As with our species data, open data sources were used where possible to ensure that our data remained in line with FAIR data practices (Wilkinson et al., 2016).

**Table 1:**
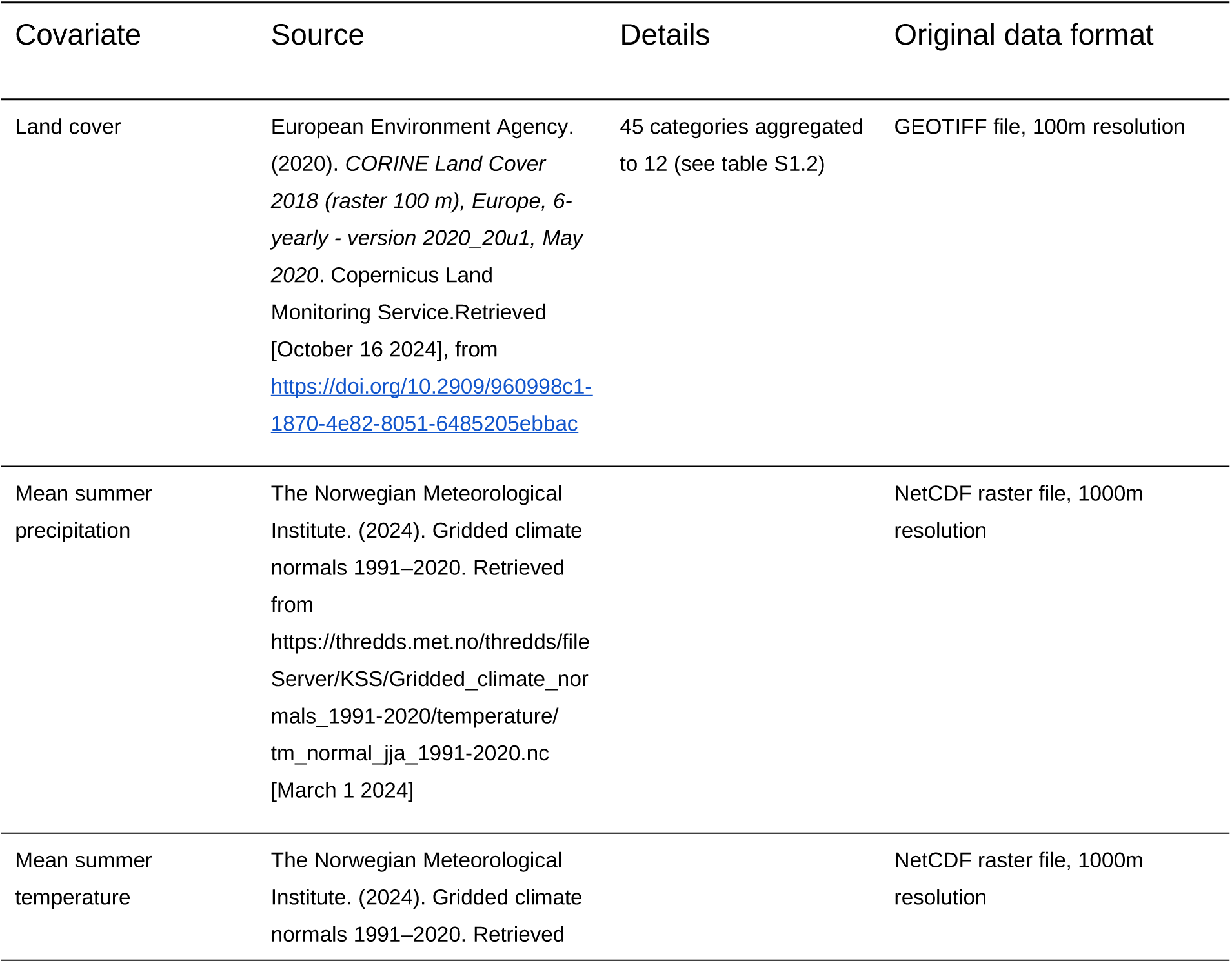
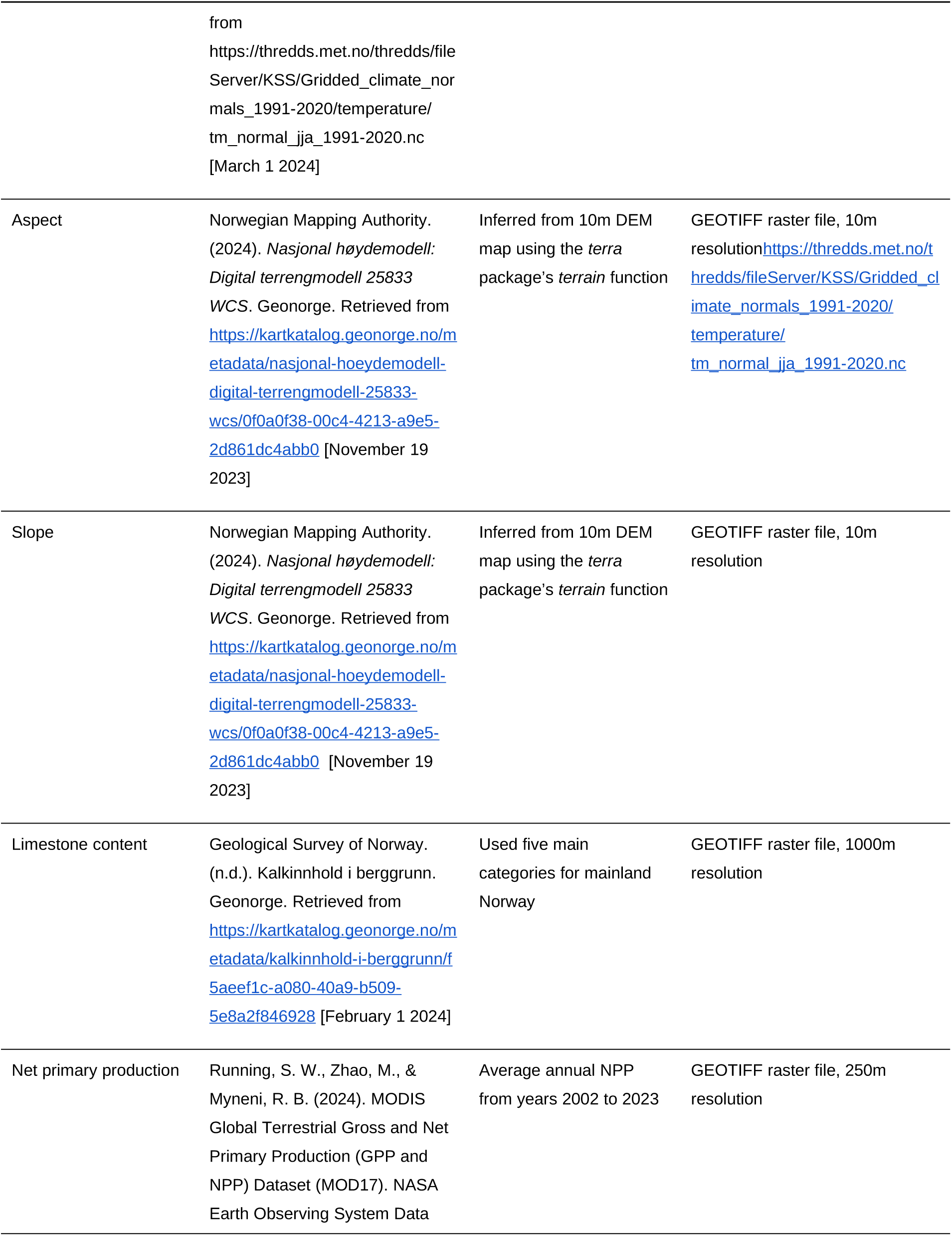
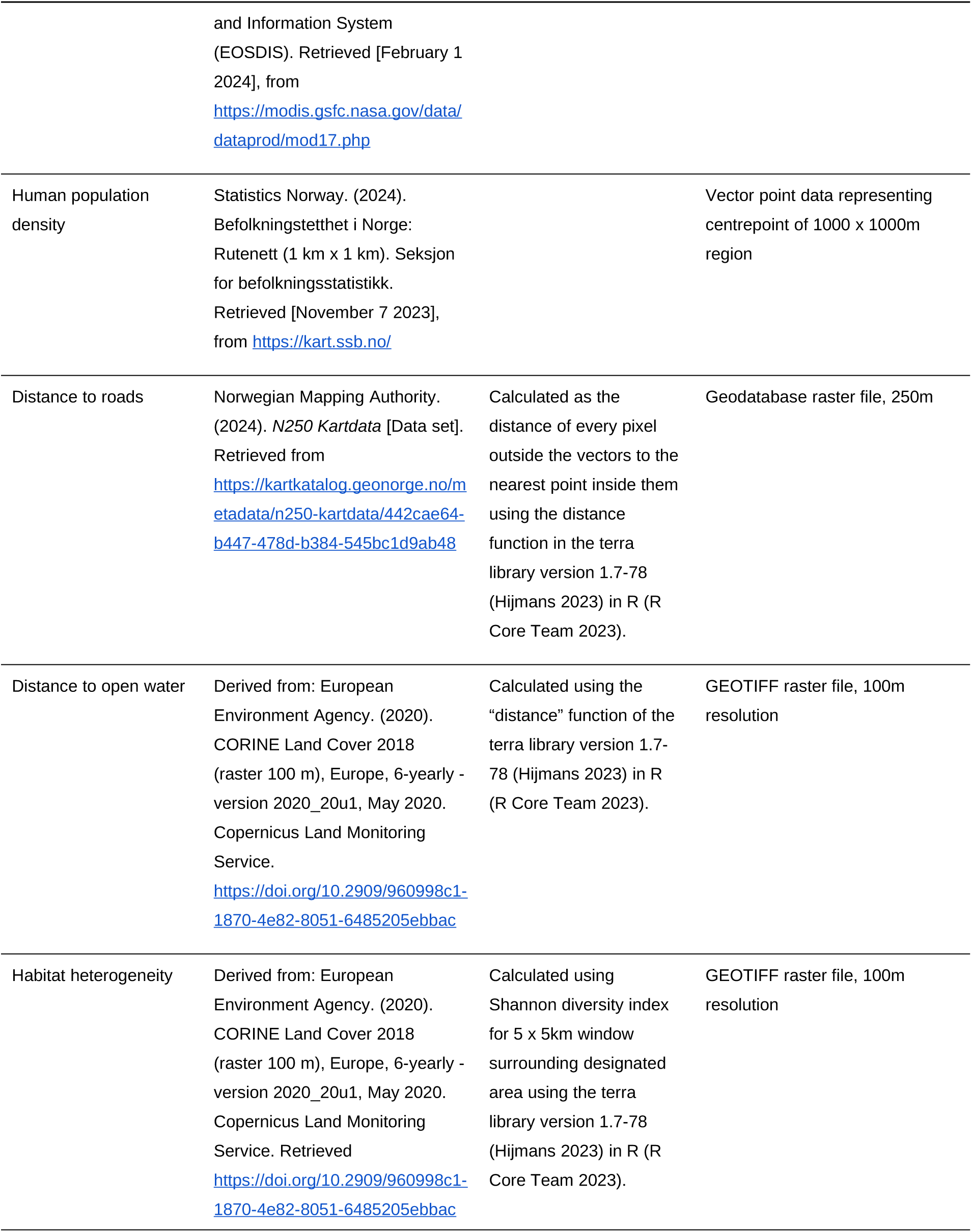
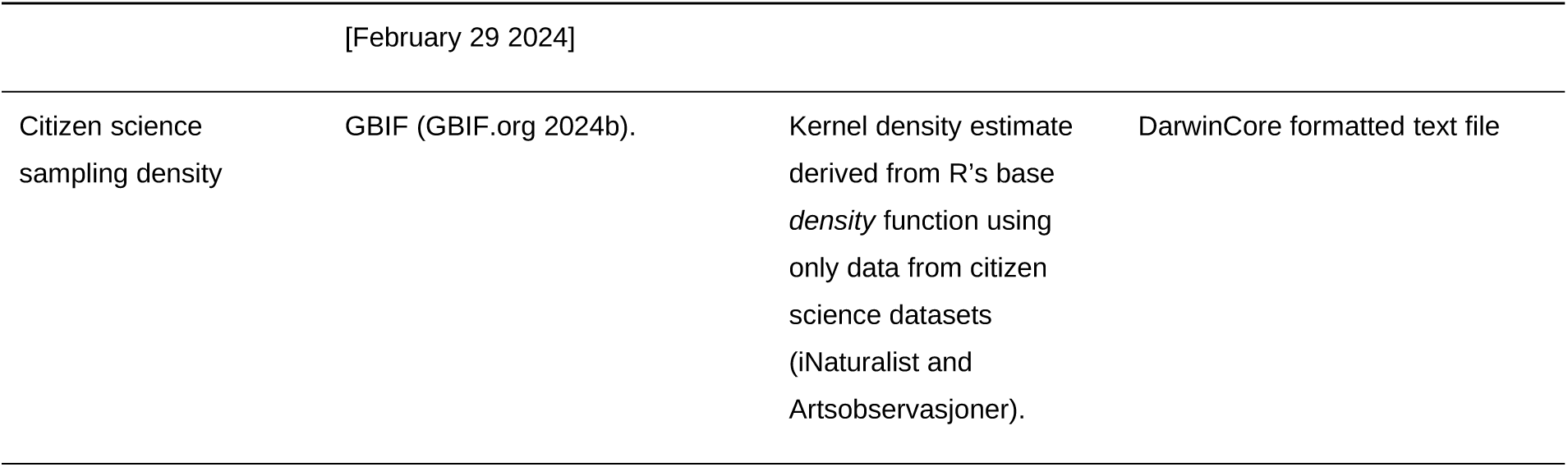
Description of environmental covariates used in integrated species distribution models estimating species richness in vascular plants across Norway.

All environmental covariates were projected to a common UTM coordinate reference system (zone 33 North, in metres) and a resolution of 500 by 500 metres. This resolution was chosen as it represented a reasonable balance between the resolution of the chosen environmental covariates, it allowed us to keep a large number of records by expanding the threshold for coordinate uncertainty to 250 metres, and was feasible to compute on the chosen infrastructure. Bilinear interpolation was used to estimate values for continuous covariates, while for categorical covariates, the most frequent category (i.e., mode) within the 500 by 500 metre cell was used. After cropping to Norwegian land borders and standardising the resolution, all continuous covariates were mean-centred and scaled by their standard deviation. Finally, we merged similar categories in the discrete land cover covariate (originally defined by 45 categories) into 12 representative categories in order to avoid an unnecessarily high number of model parameters and eliminate many categories which are not present in mainland Norway (see supplementary table S1.2).

To prevent overfitting and collinearity of variables, several of our initial environmental variables were removed due to correlated effects on species distributions. Table 1 lists the final model covariates.

### Species distribution modelling

For this framework we use integrated species distribution models (iSDMs). iSDMs combine datasets obtained from different sampling methodologies in a single model, allowing us to account for the unique biases and make use of the individual strengths within each dataset (Mostert et al. 2024a). We use point-process methods, because they are flexible when working with several data types and can deal with change of support and mis-alignment problems (Pacifici et al. 2019).

The approach builds a hierarchical state-space model, combining an underlying process model for the actual, unobserved species distribution (latent state) with multiple observation sub-models for each dataset, describing how each dataset was produced. Using this formulation, the underlying latent state and its associated parameters are shared across the different observation models, accounting for the effects of environmental parameters as well as spatial autocorrelation on species distributions. However, each dataset may also have its own parameters used to account for the way that it was collected and the biases inherent in each.

We model the true species distribution model as a log-Gaussian cox process (LGCP). This model is an inhomogeneous Poisson point-process model characterized by its spatially-varying intensity function *λ _j_* (*s*)=exp (*η_j_* (*s*)), describing the expected number of occurrences for species *j* found at any location *s* per unit area. On the log scale, this intensity function is given by:

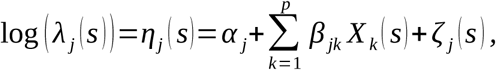

Where: *α_j_* is a species-specific intercept term, *β_jk_* is the regression coefficient for the *k^th^* environmental predictor for species *j* and *ζ* (*s*) is a species-specific spatially-varying zero-mean Gaussian random field characterized by a Matérn correlation function.

Given that estimating species-specific spatial effects is computationally difficult, we assumed that the species had different spatial effects, but shared the same hyperparameters of the model.

To account for the individual characteristics of each dataset, we use observation models, which provide a statistical description of the data collection protocol for a given dataset. For the detection/non-detection datasets, we assume a Bernoulli model. To integrate these data in the Poisson process model, we modelled the likelihood that at least one observation of species *j* at some location *s* (*N _j_*(*s*)>0) was found using the complementary log-log (cloglog: *f* (*x*)=log (*−*log (1 *−x*)) link function. Using this formulation, we may express the detection/non-detection as a function of the intensity function for our process function, denoted by:

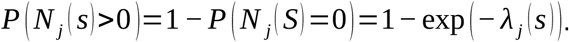

We assume that our presence-only data are prone to common sampling biases, and that these biases can be related to covariates related to accessibility, in this case represented by distance to roads and citizen science sampling intensity. As a result, we model these data as a thinned version of our Poisson point-process model. The thinning probability of the intensity function is defined by *γ* (*s*), which we can assume to be dependent on a vector of sampling covariates *Ω* (*s*) as well as a sampling bias field (Simmonds et al. 2020). Assuming this model, the thinned intensity is given as:

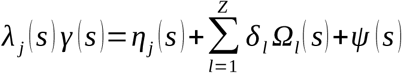

Where: *δ_l_* represents the regression coefficient for the *_l_^th^* sampling covariate, and *ψ* (*s*) is a spatial field. This model can be fitted to the data with the *PointedSDMs* R package (Mostert & O’Hara 2023). Here we use the *intSDM* R package (Mostert et al., 2024b), a wrapper package for *PointedSDMs* which is specifically designed for efficiently estimating iSDMs within a point-process framework using the integrated nested Laplace approximation (INLA) methodology (Rue et al., 2009), which is a method for approximate Bayesian inference for Latent Gaussian models. The package does so by building wrapper functions around the R package *inlabru* (Bachl et al., 2019), which is designed to facilitate spatial modelling using INLA.

Regional sampling density was calculated using R’s base *density* function. All presence records were first converted into a point pattern dataset using the *spatstat* package (Baddeley et al., 2013). The *density* function was then applied with a smoothing coefficient of 0.04. The resulting metric was then log-transformed to better visualise variations in regions of the country found further from cities, where heavy citizen science sampling skew visualisations of sampling density.

The models were fitted to all species with more than 50 presence observations. In order to maximise the number of species modelled to match up the memory available on the server, we broke up each group into segments of nine or ten species, with one additional common species well-surveyed in the representative dataset assigned to each segment. This was done so that the common species would provide information about the sampling processes that could be used to inform the models for the rarer species.

The models were fitted using HPC services available through Sigma2 - the National Infrastructure for High-Performance Computing and Data Storage in Norway. Using the 178 gigabyte memory with 20 CPUs, each segment was run to either generate the model fit or predictions of species intensity and spatial bias for the segment. The average CPU time used per vascular plant segment model was 190 hours, which is computationally efficient given the number of species included per segment and the model complexity.

### Post-processing

The output of the Poisson point process model is an intensity function, which describes the expected number of a given species per unit of area. For results to be management-relevant the actual likelihood of occurrence of a site/pixel needs to be calculated. As such, we estimate probabilities of occurrence by generating 100 samples of the model parameters from the posterior and using these to calculate probabilities of occurrence given the covariate values obtained at each pixel on the map of our study area. We assume that our model estimates from the spatially representative ANO dataset correctly predict occurrence throughout the study area, and then we scale the estimate of the final iSDM prediction using the study area of this representative dataset.

Likelihood of occurrence was obtained by using the inverse of the cloglog link function of the linear predictor – expressing the likelihood that at least one occurrence of a species occurs in a given pixel. Mathematically, the expected occupancy of species *j* for some sampling area area *A* given the intensity function *λ* is given by:

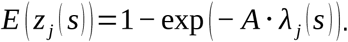

Species richness and its associated estimates of uncertainty were then estimated by summing up the individual mean for likelihood of occurrence for each species at each pixel. This was then scaled to between 0 and 1 to make it clear that the metric given was a relative term and not a direct estimation of the number of species present at each pixel. The standard error was calculated by taking the square root of the summed squared standard errors for each pixel. As we did not want known alien species included in our species richness estimates, species listed on the Alien Species List created by the Norwegian Species Database (Artsdatabanken, 2023) were removed from the summed estimates.

### Result Validation

Result validation was carried out using the Vascular Plants Field Notes Dataset published by the NTNU University Museum (NTNU, 2025). The data was collected from the late 1950s until 2006 using a standardised checklist to survey a designated polygon for all species on the checklist. Each of these surveys comprises a sampling event, with species subsequently marked as present when found. Polygons ranged in size from 0.01 to 48 square kilometres, although 82% of sampling events sampled an area of one square kilometre (±0.05 kilometres squared). All sampling events whereby the relevant polygon was larger than 0.5 kilometres squared had a coordinate uncertainty of more than 250 metres and were excluded from use in model estimation. These were instead used in result validation. Only sampling events recorded after the year 2000 were used to ensure comparability of results.

For the resulting subset of sampling events which surveyed polygons with a total area of more than one and less than ten square kilometres, we obtained observed richness by tallying all species found in each sampling event. To obtain estimated richness from our models, we extracted the maximum occurrence likelihood from all pixels that intersected with the given polygon. Resulting probabilities were then tallied to give an estimated richness. Estimated and observed richnesses were then compared using a standard linear model with observed richness as the explanatory variable and estimated richness as the response variable. Size category was also used as an extra factorial variable, separating polygons under two kilometres in area from the rest in order to account for variation in sampling completion that is likely to stem from an increase in sampling area. These results were compared to the results produced by Olsen et al. (2018), which used three presence-only datasets and as such did not employ an integrated Species Distribution Modelling approach. Comparisons for model fit were made using the R-squared metrics from both models.

## Results

### Overall workflow performance

The data acquisition section of the workflow resulted in 41 datasets, of which 28 were classified as occurrence only datasets (supplementary table S2). After the remaining 13 were subsequently processed, the resulting combination of occurrence only and presence-absence data yielded 5,977,377 records for 2880 species of vascular plants. From these we removed 1250 species because they had less than 50 records, leaving 1630 species. From this list a further 412 species were removed, as they were on the Norwegian Alien Species List (Artsdatabanken, 2023), leaving us with 1218 species. These included 147 threatened species and 178 species on the list of species of national responsibility.

### Species Richness and Distributions

The workflow produces maps of the spatial distribution of a subset of vascular plant biodiversity across Norway (Figure 2a), along with a map of the uncertainty (Figure 2b). Species richness was estimated to be distinctly lower in the northernmost and south-western regions of the country. The workflow also produces sampling density maps (Figure 2c) which show distinctly higher sampling levels in the regions surrounding heavily populated Norwegian cities in the south-east and middle of the country. There is still a notable correlation between sampling intensity and estimated species richness, though given the high correlation between human settlement and areas naturally high in biodiversity throughout Norway this is to be expected. There is substantial variation in estimated richness both inside and outside of regions with high sampling density, indicating progress in compensating for sampling bias. The species richness statistics are useful for identifying where biodiversity is highly concentrated, both at a national and regional level, with areas of high richness on a more regional level still able to be pinpointed when they are relatively low in richness on a national scale. The variation shown in uncertainty does not show the same level of regional variation as species richness, with more fine-scale variation present along the whole gradient of Norway. Uncertainty is lower both in areas with high sampling and in areas with very low species richness, most notably around the larger cities and the south-west coast. Outside of these regions the spatial mesh utilised drives uncertainty, with higher uncertainty at mesh vertices, indicating a need for better data in more remote areas and further development of spatial field implementation.

**Figure 2:**
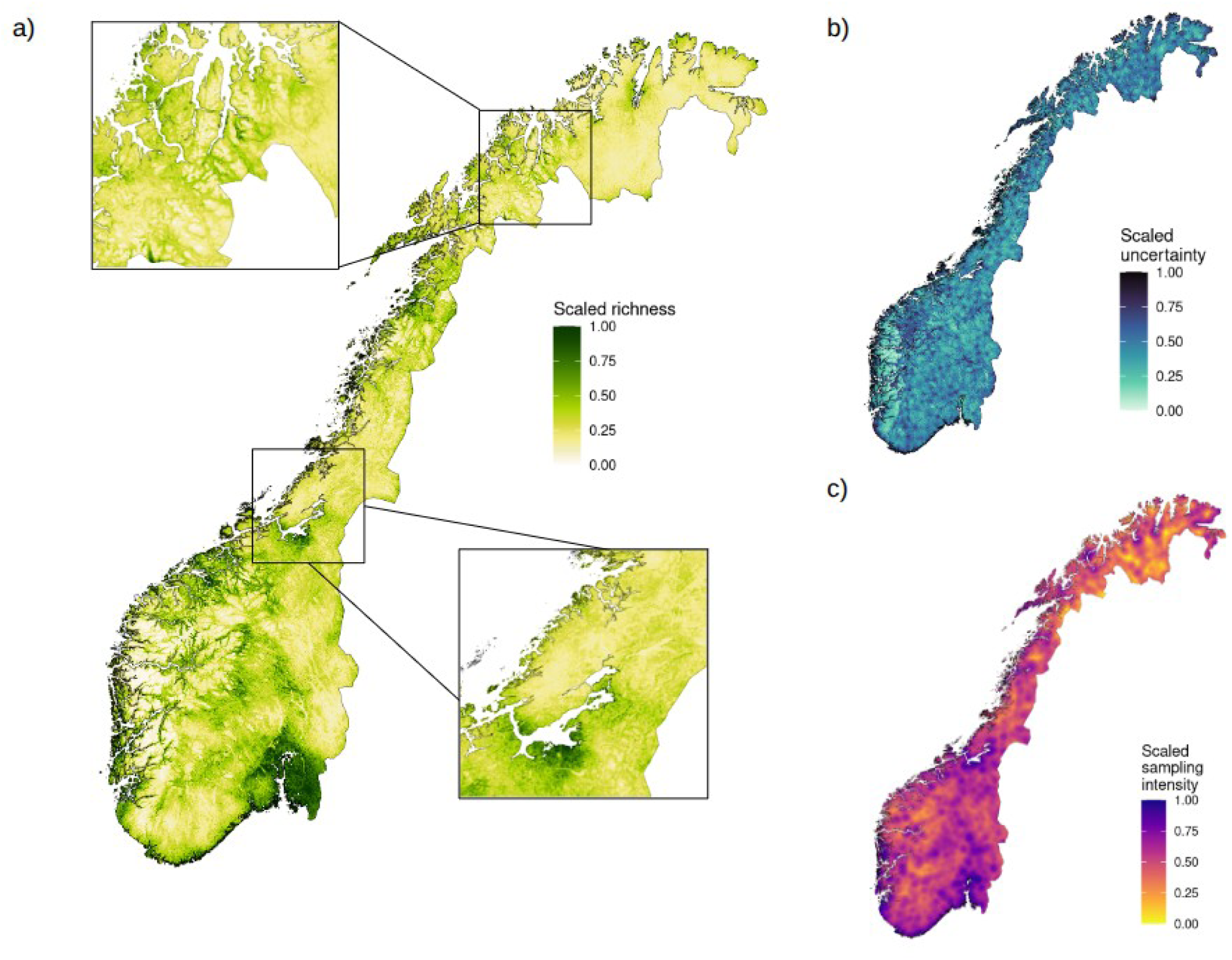
Maps of a) scaled species richness estimates across Norway, b) uncertainty for richness estimates, and c) sampling density across Norway for vascular plants. Results generated from iSDM-based workflow which incorporates open data sources. Each metric has been scaled to between 0 and 1, with 1 indicating that the metric is at its highest for Norway at this point. Furthermore, uncertainty has been truncated to between the 10th and 90th percentile of its initial values before scaling, in order to better visualise variation across the region.

Additionally, by predicting for selection of species of interest it is possible to show the same metrics for distinct groups of scientific or cultural importance. Here we have estimated richness for species classed as threatened in Norway (Artsdatabanken, 2021) (Figure 3a-c), which show similar trends to all species, and a more exaggerated difference in sampling density when moving from heavily populated to less populated areas.

**Figure 3:**
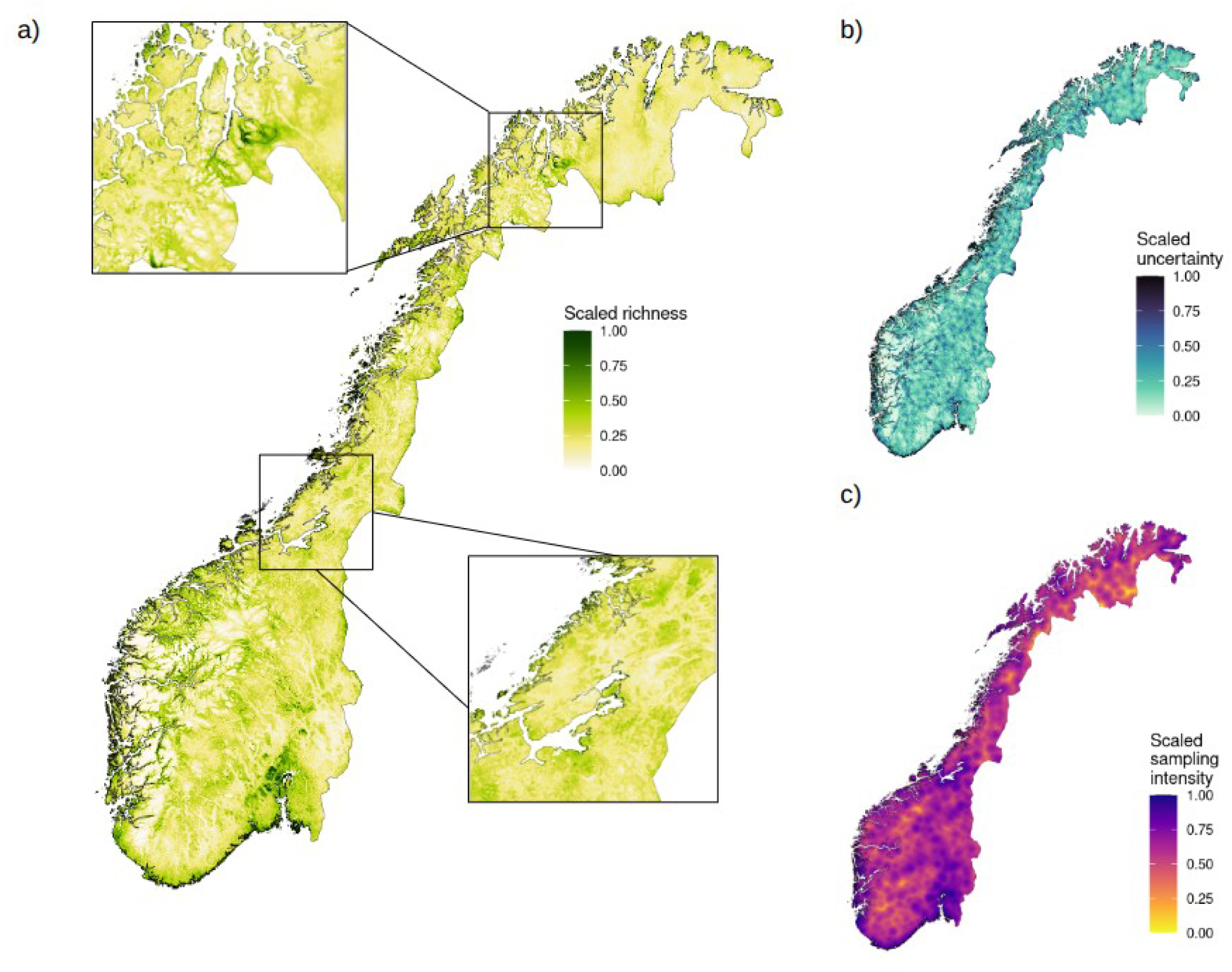
Maps of a) scaled species richness estimates across Norway, b) uncertainty for richness estimates, and c) sampling density across Norway for threatened vascular plants. Results generated from iSDM-based workflow which incorporates open data sources. Each metric has been scaled to between 0 and 1, with 1 indicating that the metric is at its highest for Norway at this point. Furthermore, uncertainty has been truncated to between the 10th and 90th percentile of its initial values before scaling, in order to better visualise variation across the region.

As well as maps, other metrics of interest can be calculated. For example, the distribution of effects of ecosystem type across species can be presented (figure S1), which suggests that, not surprisingly, plant species richness tends to be higher in grasslands than on sparsely vegetated lands.

Field validation results showed a general correlation between estimated and observed species richness (p < 0.05), albeit with a large amount of variation (R^2^ = 0.31) (figure 4a). Explanatory power increased when compared to results from Olsen et al. (2018) (R^2^ = 0.26, figure 4b).

**Figure 4:**
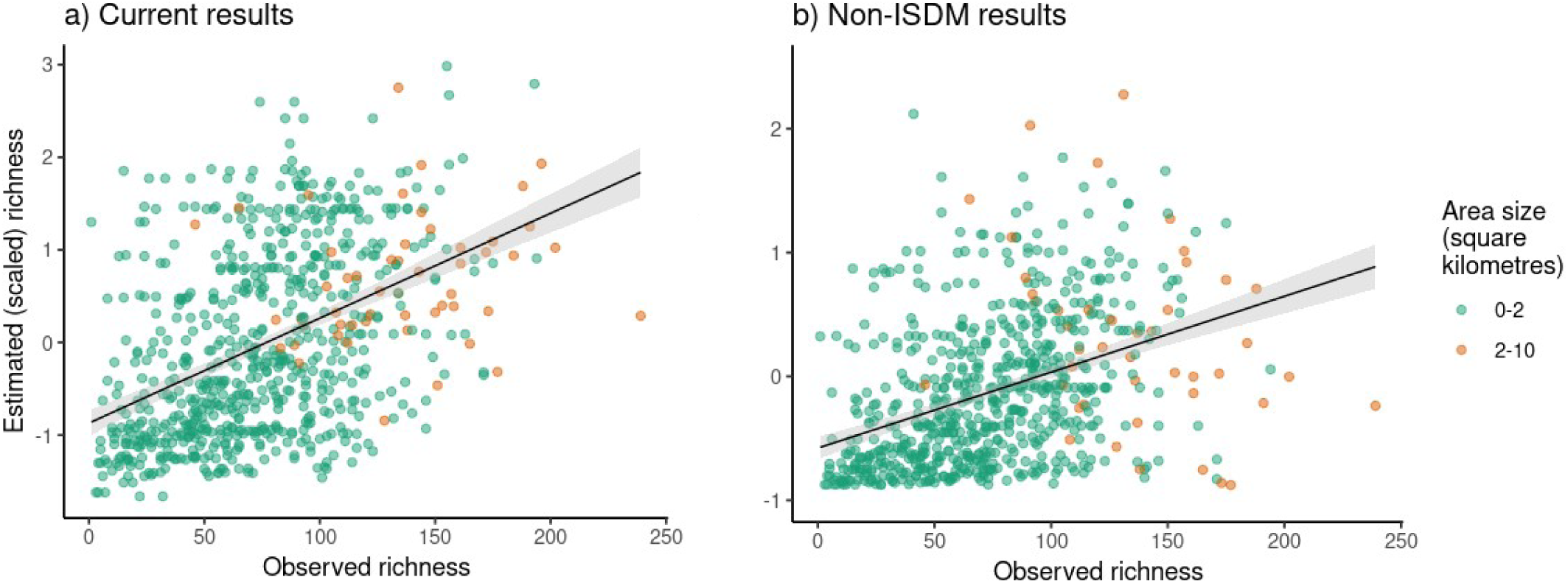
Comparison of estimated richness generated by (a) the current point-process based iSDM pipeline and (b) previous non integrated models. Richness estimates generated by aggregating likely presences. Observed richness calculated using sampling events in a structured dataset of vascular plant Field Notes (NTNU, 2018).

## Discussion

The workflow presented here represents a significant advance in the estimation of large-scale biodiversity patterns. By facilitating the integration of disparate datasets and addressing biases in collection methods, it provides a robust framework for biodiversity modeling that can be used to process a large number of species. With outputs that include individual species occurrence probabilities, species richness estimates, uncertainty metrics and sampling effort, the workflow offers a comprehensive suite of tools to tackle some of the most pressing challenges in applied biodiversity science.

The approach here prioritizes breadth over depth, so that individual species maps will be less precise than if models for each species were tailored to that species, for example by customizing covariance and model prior selection. As such, decisions made with the conservation of individual species or particular sub-groups of species may not be supported by this pipeline. But the gain is a holistic perspective on spatial patterns of biodiversity. This broader focus is particularly valuable for conservation planning, enabling the identification of regions of high biodiversity significance and informing strategic decision-making.

Although the models for individual species are not optimal, one advantage of modelling a large number of species is that we can correct for sampling biases and detectability by leveraging information across species (Isaac & Pocock 2015). In the future, this should help in the development of genuine community models.

As highlighted above, several key areas require improvement to enhance the workflow’s effectiveness, spanning ecological, statistical, and computational dimensions.

### Improving the Workflow

The development of a modelling workflow must be updated regularly in order to take advantage of advances in data, technology and methodology. The current software needs to be extended to be able to import a wider range of species observation and environmental data, and also work with more model fitting packages than *PointedSDMs* (Mostert & O’Hara 2023). After the modelling, the returned objects also need to have a common format, so that the post-processing can be run regardless of model used.

Improving covariate modeling represents a critical avenue for advancing species distribution models. While the current selection of covariates is guided by expert judgment, it lacks systematic evaluation of their utility for predicting species distributions. Future efforts should focus on identifying optimal covariates for modeling effort and exploring alternative approaches, such as spatial field-based methods. Temporal variability in environmental effects also warrants attention, as changing conditions can significantly influence species distributions. In the longer term, spatio-temporal models offer the potential to capture these dynamics more accurately, albeit with increased computational demands (Seaton et al. 2024). Additionally, incorporating phenology (e.g., seasonal availability of species) and community-level models (taking into account species associations) could further enhance ecological relevance.

Expanding model validation and diagnostic methods is equally important to ensure reliability and to refine both the statistical and ecological components of the workflow. The observation effort model also requires improvement, particularly in addressing sampling biases. This could involve incorporating more relevant covariates or employing spatial random fields. A robust, generalizable framework for modeling these biases is essential, particularly for effectively utilizing citizen science data.

### A Call for Better Data

A key limitation of current biodiversity modeling efforts in general, including the workflow presented here, is the lack of sufficiently structured and standardized data. While we have demonstrated that the integration of disparate datasets allows for significant advances in addressing large-scale biodiversity patterns, more readily available metadata would lead to better data integration and subsequent model accuracy. These issues limit the scalability and adaptability of our results to diverse ecological and management contexts.

One clear limitation of our results produced by a lack of data is the remaining correlation between sampling intensity and estimated species richness. Even though human settlements are typically found in high-productivity areas, and semi-natural areas created by human activity often result in high vascular plant richness (Austrheim, 2002; Pautasso, 2007), a lack of sampling in more remote areas can still restrict our accuracy. While our models account for habitat suitability, our use of a spatial field can restrict the estimated distribution of species if recorded observations of the species are very localised.

Structured and well-documented data are foundational to resolving these challenges. Datasets with precisely recorded presences and absences or quantitative data are crucial, as is the recording of sampling effort, and increased availability of such datasets allow for more precise bias correction. Additionally, while metadata analysis is currently a time-consuming manual process, this process could be streamlined through the development of robust criteria for this analysis and subsequent integration into an AI or machine-learning framework, both of which are well-suited for scaling up of traditional data analysis (Ullah et al., 2024). Measures like those proposed in Moersberger et al. (2024), including enhancing cooperation between data collectors and increasing stakeholder engagement, can help mobilise data and increase model accuracy.

The above suggestions highlight the need for a coordinated effort to mobilize and integrate relevant datasets. Mapping and monitoring data from governmental sources, such as those managed by the Norwegian Environmental Agency, represent a valuable but often untapped resource. Efforts to clean, standardize, and make these datasets openly available according to FAIR principles would significantly enhance the breadth and depth of biodiversity modeling. Similarly, the incorporation of more structured citizen science data, if properly curated, offers an opportunity to complement traditional datasets and address spatial or taxonomic biases. Mondain-Monval et al. showed that directing citizen scientists to previously undersurveyed areas can improve model performance (2024). Additionally, increased availability of environmental observations following occurrences of organisms can facilitate the identification of ecologically meaningful covariates and improve subsequent estimates. The development and adoption of community-driven standards like the Darwin Core Archive for data collection, storage, and sharing will be critical to achieving this goal. This can be achieved through outreach programs, but also through the implementation of maps like those produced here to demonstrate the usefulness of open data.

By addressing the current limitations in data availability and standardization, future workflows can provide even more precise, reliable, and actionable insights for ecological research and conservation planning. Achieving this will require collaboration across scientific, governmental, and citizen science communities to build the comprehensive, structured data resources needed to support the next generation of biodiversity modeling.

## Author contributions

SWP, KPA, PSM and RRT wrote code necessary for result production. KPA and SWP ran models and produced end product data. SWP, IH and RRT sourced input data for models. JT, JC, RBO and AGF provided guidance in model construction during project. JAG and IH supervised and reported on field data collection. All authors contributed to the writing of the paper.

## Data availability statement

All code necessary is available at the gjærvoll/BioDivMapping repository, linked here: https://doi.org/10.5281/zenodo.19328599. The code includes data sources for all environmental variables. The reference section provides DOIs which link to the necessary species occurrence data.

## Acknowledgements

Ron Togunov was supported by a grant from the Research Council of Norway (project: GreenPlan, project no. 326979). No fieldwork permits were required.

