## supplementaryInformation for "Addressing Data Fragmentation in Biodiversity: A Workflow for integrated Species Distribution Models"

**Content**

[**S1. Choice of environmental covariates	1**](https://docs.google.com/document/d/1suzkrOQR1LJBDnTMFt8ksq9RoZ3Wh0aX3drSItnb8EY/edit?tab=t.0#heading=h.d9499vf47yib)

[**S2. Observations per dataset for vascular plants**](https://docs.google.com/document/d/1suzkrOQR1LJBDnTMFt8ksq9RoZ3Wh0aX3drSItnb8EY/edit?tab=t.0#heading=h.5942oddi6fej)	**6**

[**S3. Field Validation of Modelling Outputs**](https://docs.google.com/document/d/1suzkrOQR1LJBDnTMFt8ksq9RoZ3Wh0aX3drSItnb8EY/edit?tab=t.0#heading=h.9t5m16e3prgo)	**9**

[S3.1 Data collection](https://docs.google.com/document/d/1suzkrOQR1LJBDnTMFt8ksq9RoZ3Wh0aX3drSItnb8EY/edit?tab=t.0#heading=h.fopg0jud47jz)	9

[S3.2 Result comparison](https://docs.google.com/document/d/1suzkrOQR1LJBDnTMFt8ksq9RoZ3Wh0aX3drSItnb8EY/edit?tab=t.0#heading=h.4hkcqy5rfv46)	10

[**References**](https://docs.google.com/document/d/1suzkrOQR1LJBDnTMFt8ksq9RoZ3Wh0aX3drSItnb8EY/edit?tab=t.0#heading=h.w7wv4rqicmh7)	**11**

### S1. Choice of environmental covariates

| *Table S1.1: List of environmental covariates considered for use in our integrated Species Distribution Model workflow.* | | | |
| --- | --- | --- | --- |
| Covariate | Source | Details | Original data format |
| Land cover | European Environment Agency. (2020). *CORINE Land Cover 2018 (raster 100 m), Europe, 6-yearly - version 2020_20u1, May 2020*. Copernicus Land Monitoring Service.Retrieved [October 16 2024], from <https://doi.org/10.2909/960998c1-1870-4e82-8051-6485205ebbac> | 45 categories aggregated to 12 (see table S1b) | GEOTIFF file, 100m resolution |
| Mean summer precipitation | Meteorologisk institutt. (2024). Gridded climate normals 1991–2020. Retrieved from https://thredds.met.no/thredds/fileServer/KSS/Gridded_climate_normals_1991-2020/temperature/tm_normal_jja_1991-2020.nc [March 1 2024] |  | NetCDF raster file, 1000m resolution |
| Mean summer temperature | Meteorologisk institutt. (2024). Gridded climate normals 1991–2020. Retrieved from https://thredds.met.no/thredds/fileServer/KSS/Gridded_climate_normals_1991-2020/temperature/tm_normal_jja_1991-2020.nc [March 1 2024] |  | NetCDF raster file, 1000m resolution |
| Aspect | Norwegian Mapping Authority. (2024). *Nasjonal høydemodell: Digital terrengmodell 25833 WCS*. Geonorge. Retrieved from <https://kartkatalog.geonorge.no/metadata/nasjonal-hoeydemodell-digital-terrengmodell-25833-wcs/0f0a0f38-00c4-4213-a9e5-2d861dc4abb0> [November 19 2023] | Inferred from 10m DEM map using the *terra* package’s *terrain* function | GEOTIFF raster file, 10m resolution<https://thredds.met.no/thredds/fileServer/KSS/Gridded_climate_normals_1991-2020/temperature/tm_normal_jja_1991-2020.nc> |
| Slope | Norwegian Mapping Authority. (2024). *Nasjonal høydemodell: Digital terrengmodell 25833 WCS*. Geonorge. Retrieved from <https://kartkatalog.geonorge.no/metadata/nasjonal-hoeydemodell-digital-terrengmodell-25833-wcs/0f0a0f38-00c4-4213-a9e5-2d861dc4abb0> [November 19 2023] | Inferred from 10m DEM map using the *terra* package’s *terrain* function | GEOTIFF raster file, 10m resolution |
| Limestone content | Norges geologiske undersøkelse. (n.d.). Kalkinnhold i berggrunn. Geonorge. Retrieved from <https://kartkatalog.geonorge.no/metadata/kalkinnhold-i-berggrunn/f5aeef1c-a080-40a9-b509-5e8a2f846928> [February 1 2024] | Used five main categories for mainland Norway | GEOTIFF raster file, 1000m resolution |
| Net primary production | Running, S. W., Zhao, M., & Myneni, R. B. (2024). MODIS Global Terrestrial Gross and Net Primary Production (GPP and NPP) Dataset (MOD17). NASA Earth Observing System Data and Information System (EOSDIS). Retrieved [February 1 2024], from <https://modis.gsfc.nasa.gov/data/dataprod/mod17.php> | Average annual NPP from years 2002 to 2023 | GEOTIFF raster file, 250m resolution |
| Human population density | Statistisk sentralbyrå. (2024). Befolkningstetthet i Norge: Rutenett (1 km x 1 km). Seksjon for befolkningsstatistikk. Retrieved [November 7 2023], from <https://kart.ssb.no/> |  | Vector point data representing centrepoint of 1000 x 1000m region |
| Distance to roads | Statens kartverk. (2024). *N250 Kartdata* [Data set]. Retrieved from <https://kartkatalog.geonorge.no/metadata/n250-kartdata/442cae64-b447-478d-b384-545bc1d9ab48> | Calculated as the distance of every pixel outside the vectors to the nearest point inside them using the distance function in the terra library version 1.7-78 (Hijmans 2023) in R (R Core Team 2023). | Geodatabase raster file, 250m |
| Distance to open water | Derived from: European Environment Agency. (2020). CORINE Land Cover 2018 (raster 100 m), Europe, 6-yearly - version 2020_20u1, May 2020. Copernicus Land Monitoring Service. <https://doi.org/10.2909/960998c1-1870-4e82-8051-6485205ebbac> | Calculated using the “distance” function of the terra library version 1.7-78 (Hijmans 2023) in R (R Core Team 2023). | GEOTIFF raster file, 100m resolution |
| Habitat heterogeneity | Derived from: European Environment Agency. (2020). CORINE Land Cover 2018 (raster 100 m), Europe, 6-yearly - version 2020_20u1, May 2020. Copernicus Land Monitoring Service. Retrieved  <https://doi.org/10.2909/960998c1-1870-4e82-8051-6485205ebbac> [February 29 2024] | Calculated using Shannon diversity index for 5 x 5km window surrounding designated area using the terra library version 1.7-78 (Hijmans 2023) in R (R Core Team 2023). | GEOTIFF raster file, 100m resolution |
| Extreme temperature events | Meteorologisk institutt. (2024). Gridded climate normals 1991–2020: Summer temperature (JJA). Retrieved from https://thredds.met.no/thredds/fileServer/KSS/Gridded_climate_normals_1991-2020/temperature/tm_normal_jja_1991-2020.nc | l | NetCDF raster file, 1000m resolution |
| Elevation | Norwegian Mapping Authority. (2024). *Nasjonal høydemodell: Digital terrengmodell 25833 WCS*. Geonorge. Retrieved from <https://kartkatalog.geonorge.no/metadata/nasjonal-hoeydemodell-digital-terrengmodell-25833-wcs/0f0a0f38-00c4-4213-a9e5-2d861dc4abb0> [Novemeber 19 2023] |  | GEOTIFF raster file, 10m resolution |
| Soil coarseness | Norges geologiske undersøkelse. (2024). *Løsmasser forenklet kart N1000*. Retrieved from <https://kartkatalog.geonorge.no/metadata/loesmasser-forenklet-kart-n1000/fd37fe85-ab6d-4e9e-8ca8-f09fffdba86d> [February 1 2024] |  | Shapefile raster, 1000m resolution |
| Maximum NDVI | Didan, K. (2024). MOD13A2: MODIS Normalized Difference Vegetation Index (NDVI). NASA EOSDIS Land Processes DAAC. Retrieved from <https://lpdaac.usgs.gov/products/mod13a2v006/> [February 1 2024] |  | GEOTIFF raster file, 250m resolution |
| Building density | Statistisk sentralbyrå. (2024). *Bygningsmasse i Norge: Rutenett (1 km x 1 km)*. Seksjon for eiendoms-, areal- og primærnæringsstatistikk. Retrieved from <https://kart.ssb.no/> [November 19 2023] |  | Vector point data representing centrepoint of 1000 x 1000m region |
| Duration of snow cover | Karger, D. N., Lange, S., Hari, C., Reyer, C. P. O., & Zimmermann, N. E. (2022). *CHELSA-W5E5 v1.0: W5E5 v1.0 downscaled with CHELSA v2.0*. ISIMIP. Retrieved from: https://doi.org/10.48364/ISIMIP.836809.3 [January 16 2024] |  | GEOTIFF raster file, 30 arcsec spatial resolution |
| Distance to forest line | Bryn, A., Potthoff, K. Elevational treeline and forest line dynamics in Norwegian mountain areas – a review. Landscape Ecol 33, Retrieved 1225–1245 (2018). <https://doi.org/10.1007/s10980-018-0670-8> [February 1 2024] |  | Raster file, 500m resolution |

| *Table S1.2: Simplified ecologically relevant categories applied to CORINE data in order to reduce computational time and increase ecological relevance when modelling their effects.* | |
| --- | --- |
| Original CORINE Category | Merged Category |
| Continuous urban fabric | Built up area |
| Discontinuous urban fabric | Built up area |
| Industrial or commercial units | Built up area |
| Road and rail networks and associated land | Built up area |
| Port areas | Built up area |
| Airports | Built up area |
| Mineral extraction sites | Built up area |
| Dump sites | Built up area |
| Construction sites | Built up area |
| Green urban areas | Constructed green space |
| Sport and leisure facilities | Built up area |
| Non-irrigated arable land | Constructed green space |
| Permanently irrigated land | Constructed green space |
| Rice fields | Agro-forestry areas |
| Vineyards | Agro-forestry areas |
| Fruit trees and berry plantations | Agro-forestry areas |
| Olive groves | Agro-forestry areas |
| Pastures | Constructed green space |
| Annual crops associated with permanent crops | Constructed green space |
| Complex cultivation patterns | Agro-forestry areas |
| Land principally occupied by agriculture with significant areas of natural vegetation | Agro-forestry areas |
| Agro-forestry areas | Agro-forestry areas |
| Broad-leaved forest | Broad-leaved forest |
| Coniferous forest | Coniferous forest |
| Mixed forest | Mixed forest |
| Natural grasslands | Natural grasslands |
| Moors and heathland | Moors and heathland |
| Sclerophyllous vegetation | Sclerophyllous vegetation |
| Transitional woodland-shrub | Transitional woodland-shrub |
| Beaches dunes sands | Beaches dunes sands |
| Bare rocks | Bare rocks |
| Sparsely vegetated areas | Sparsely vegetated areas |
| Burnt areas | Burnt areas |
| Glaciers and perpetual snow | Glaciers and perpetual snow |
| Inland marshes | Marsh/bog/fen |
| Peat bogs | Marsh/bog/fen |
| Salt marshes | Marsh/bog/fen |
| Salines | Water bodies |
| Intertidal flats | Water bodies |
| Water courses | Water bodies |
| Water bodies | Water bodies |
| Coastal lagoons | Water bodies |
| Estuaries | Water bodies |
| Sea and ocean | Water bodies |

### S2. Observations per dataset for vascular plants

| *Table S2: Number of observations per dataset for GBIF occurrence download which provided species occurrence data for integrated Species Distribution Modelling Workflow* | | |
| --- | --- | --- |
| Dataset name | # observations | Data type |
| ANOData | 8297190 | Presence-absence |
| ARKO strandeng | 1448 | Occurrence only |
| Artportalen | 4331 | Occurrence only |
| Ecofact | 511 | Occurrence only |
| Effects of vegetation clearing on vascular plants in power line clearings southeast Norway | 303534 | Presence-absence |
| Hatikka.fi observations | 178 | Occurrence only |
| Insects of the Forest-Tundra Ecotone (ForTunE) | 64 | Occurrence only |
| Jordal | 98239 | Occurrence only |
| Monitoring data of natural and man-made semi-natural meadows in and around Oslo, Norway 2018-2021 | 8332 | Presence-absence |
| Norwegian Biodiversity Information Centre - Other datasets | 140 | Occurrence only |
| Norwegian Species Observation Service | 2532797 | Occurrence only |
| Norwegian specimens stored in Herb. Klepsland (pre 2022) | 71 | Occurrence only |
| Observation.org, Nature data from around the World | 20408 | Occurrence only |
| Occurrence data from various smaller projects in Norway | 6740 | Occurrence only |
| Overvåking av semi-naturlig eng (ASO) | 89996 | Presence-absence |
| Overvåking av åpen grunnlendt kalkmark i Oslofjordområdet | 105258 | Presence-absence |
| Pl@ntNet automatically identified occurrences | 21802 | Occurrence only |
| Pl@ntNet observations | 1973 | Occurrence only |
| Stabbetorp - Floristiske registreringer 2016 | 9043 | Occurrence only |
| Vascular Plant Herbarium, Oslo (O) UiO | 86676 | Occurrence only |
| Vascular Plant Herbarium, UiB | 13419 | Occurrence only |
| Vascular Plants, Field notes, Agder naturmuseum (KMN) | 178033 | Presence-absence |
| Vascular Plants, Field notes, Oslo (O) | 29236 | Presence-absence |
| Vascular Plants, Museum of Archaeology, University of Stavanger | 957 | Occurrence only |
| Vascular Plants, Observations, Oslo (O) | 1019083 | Presence-absence |
| Vascular plant field notes, NTNU University Museum | 167660 | Presence-absence |
| Vascular plant herbarium (KMN) UiA | 31752 | Occurrence only |
| Vascular plant herbarium TRH, NTNU University Museum | 27556 | Occurrence only |
| Vascular plant herbarium, The Arctic University Museum of Norway (TROM) | 21445 | Occurrence only |
| Vascular plants in power line clearings and the nearby forest, southeast Norway | 212995 | Presence-absence |
| Vegetation data with and without experimental warming, alpine Finse 2000, 2004, 2011 | 92880 | Presence-absence |
| iNaturalist Research-grade Observations | 36683 | Occurrence only |
| naturgucker | 115 | Occurrence only |

### S3. Derived products

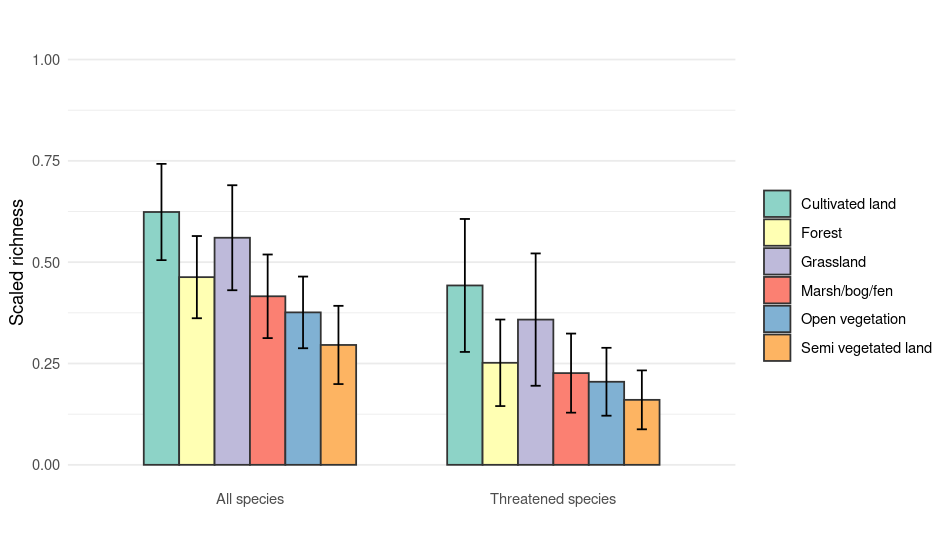

Figure S1: Comparison of relative species richness for vascular plants in Norway (both all species and threatened species) across different habitat types.
